# Insights from deconvolution of cell subtype proportions enhance the interpretation of functional genomic data

**DOI:** 10.1101/254441

**Authors:** Yu Kong, Deepa Rastogi, Cathal Seoighe, John M. Greally, Masako Suzuki

**Affiliations:** Department of Genetics and Center for Epigenomics, Albert Einstein College of Medicine, 1301 Morris Park Avenue, Bronx, New York, 10461, USA; Department of Pediatrics, Albert Einstein College of Medicine, 3415 Bainbridge Avenue, Bronx, New York, 10467, New York, USA; School of Mathematics, Statistics and Applied Mathematics, National University of Ireland Galway, University Road, Galway, H91 TK33, Ireland

## Abstract

Cell subtype proportional differences between samples significantly contribute to variation of functional genomic properties such as gene expression or DNA methylation. Current analytical approaches typically deal with cell subtype proportion influences as a nuisance variable to be eliminated. Here we demonstrate how harvesting information about cell subtype proportions from functional genomics data provides insights into the cellular events in human phenotypes. We note a striking concordance between cell subtype proportions estimated from orthogonal genome-wide assays, and demonstrate the potential for single-cell RNA-seq data to be used in tissues for which reference cell subtype functional genomic datasets are not available. Taken together, our results confirm the importance of estimating cell subtype proportions when testing a model of cellular reprogramming in human phenotypic association studies, and the value of simultaneously testing for systematic cell subtype proportional alterations as a separate phenotypic association, gaining extra insights from functional genomic studies.

## INTRODUCTION

Assays that test how the genome functions are used to understand the cellular basis for differences in phenotypes between individuals. In human disease studies, we invariably test samples that are composed of mixed populations of cell subtypes. Commonly used functional genomic assays include gene expression profiling and assays testing DNA methylation. DNA methylation can vary at a locus if it is methylated in one cell subtype but not another in the mixed sample tested, and if the proportion of these two cell subtypes differs between individuals. This was systematically demonstrated by Houseman and colleagues in studies of peripheral blood leukocytes^1^.He and others developed approaches to eliminate the influence of cell subtype proportional variation^1–4^, with the goal of eliminating an influence confounding the ability to detect changes of DNA methylation occurring in the cells studied, which have been described as cell-intrinsic changes^5^.While the same issue has been recognized to influence gene expression studies and has prompted some innovative approaches to identify cell-intrinsic changes in gene expression^6–12^, these approaches appear to be applied much less frequently in transcriptomic than in DNA methylation studies.

We have recently described our interest in understanding phenotypic associations, not only in terms of cellular reprogramming consisting of cell-intrinsic functional genomic changes, as is typically studied, but also the generation of distinctive repertoires of cell subtypes in those with a distinctive phenotype, which we have described as polycreodism^13^. As each can be distinctively informative in understanding how a phenotype developed, they could both be considered valuable insights from functional genomic studies.

We describe in this report how our re-analysis of several published studies discriminates between the separate outcomes of cell subtype changes and cell-intrinsic gene expression and DNA methylation differences associated with the phenotypes studied. We also test how gene expression and DNA methylation studies perform in their prediction of cell subtype proportions, and their concordance with each other. While we focus on studies of peripheral blood leukocytes in this evaluation, we show the potential for single cell RNA-seq to gain insights into less well characterized tissue types.These reference-based approaches are compared with the commonly-used surrogate variable analysis (SVA) approach^14,15^, testing how SVA performs to adjust for cell subtype proportion effects and other sources of variability in the data studied. We conclude that characterization of cell subtype proportions should be harvested as an outcome rather than discarded as merely a confounding influence, testing both the cellular reprogramming and polycreodism cellular models in phenotypic association studies.

## RESULTS

### Datasets used in this study

We used publicly-available datasets from the Gene Expression Omnibus (GEO), focusing on studies of peripheral blood, for which reference datasets from cell subtypes of gene expression, DNA methylation and single cell RNA-seq are better developed than for other tissues. A total of 851 peripheral blood leukocyte gene expression results and 381 DNA methylation profiles were analyzed, studying disease phenotypes involving mediation by the immune system (asthma^16,17^ and systemic lupus erythematosus (SLE)^18^), the physiological process of aging^19^, and a longitudinal study of gene expression in the same individual over 18 months^20^ (Table 1).All datasets were assessed for quality, including the elimination of samples from further analysis when there was evidence of misidentification (for example, supposedly female samples expressing genes from the Y chromosome).We describe these results in detail in the **Supplementary Note 1**.

**Table 1:** A description of the data sets used in this study.

| Study number | GEO project identifiers | Study design | Case/control | Sample numbers used in this study | Analyzed sample type |  |  |  |
| --- | --- | --- | --- | --- | --- | --- | --- | --- |
|  |  |  |  |  | Expression |  | DNA methylation |  |
| Study 1 | GSE69683 | Cross-sectional | Severe asthma/healthy | 422 | WB | Affymetrix HT HG-U133+ PM Array | N.A. |  |
| Study 2 | GSE58122 | Longitudinal | Healthy | 48 | PBMC | RNA-seq | N.A. |  |
| Study 3 | GSE40736 | Cross-sectional | Asthma/nonasthmatic | 194 | PBMC | NimbleGen Homo sapiens Expression Array | PBMC | Illumina 450k infinium |
| Study 4 | GSE82221 | Cross-sectional | SLE/healthy | 33 | PBMC | Illumina HumanHT-12 V4.0 expression beadchip | WB | Illumina 450k infinium |
| Study 5 | GSE65219, GSE58888 | Cross-sectional | Nonagenarian/young | 154 | PBMC | Illumina HumanHT-12 V4.0 expression beadchip | PBMC | Illumina 450k infinium |

SLE, systemic lupus erythematosus; N.A., not analyzed; PBMC, peripheral blood mononuclear cell; WB, whole blood; PMID, PubMed identi

### Cell subtype proportions influence gene expression results

Our first analysis was of gene expression profiles of peripheral blood leukocytes, in a study originally designed to compare individuals with severe and moderate asthma and healthy controls^16^.As variation in gene expression levels between individuals can be due to a combination of alterations of gene activity within cells (cell-intrinsic changes) as well as alterations of the proportions of cell subtypes in the sample, our goal was to understand the degree to which each mechanism was influencing the gene expression changes observed by the authors.

We took advantage of the availability of reference expression profiles for 22 different subtypes of leukocytes (LM22)^21^ and the *CIBERSORT* program, which uses a penalized multivariate regression approach to infer cell subtype proportions^12^, allowing us to estimate blood cell subtype proportions in samples from 422 individuals, both patients with severe asthma (204 females, 130 males) and healthy controls (34 females, 53 males). We then performed a principal component analysis (PCA) to estimate the contributions to gene expression variability from disease status as well as sex, smoking, and race, in addition to the influence of each estimated cell subtype proportion (Figure 1a). To measure the contribution of each covariate, we used a linear modeling approach. The principal components (PCs) of variation of the expression profiles were modeled as a linear function of cell subtype proportions. Although disease status (severe asthma) was very weakly correlated with the first two PCs of gene expression variation (PC1 (20.9% of the variance), R^2^=0.0031 and p=0.25, PC2 (10.68% of variance) R^2^=0.041 and p=2.9*10^−5^), cell subtype proportion variation showed a much greater influence on gene expression (**Supplementary Table 1**).These results indicate that for this asthma dataset, the major determinant of gene expression variation was cell subtype proportion rather than the disease phenotype.

**Figure 1.**
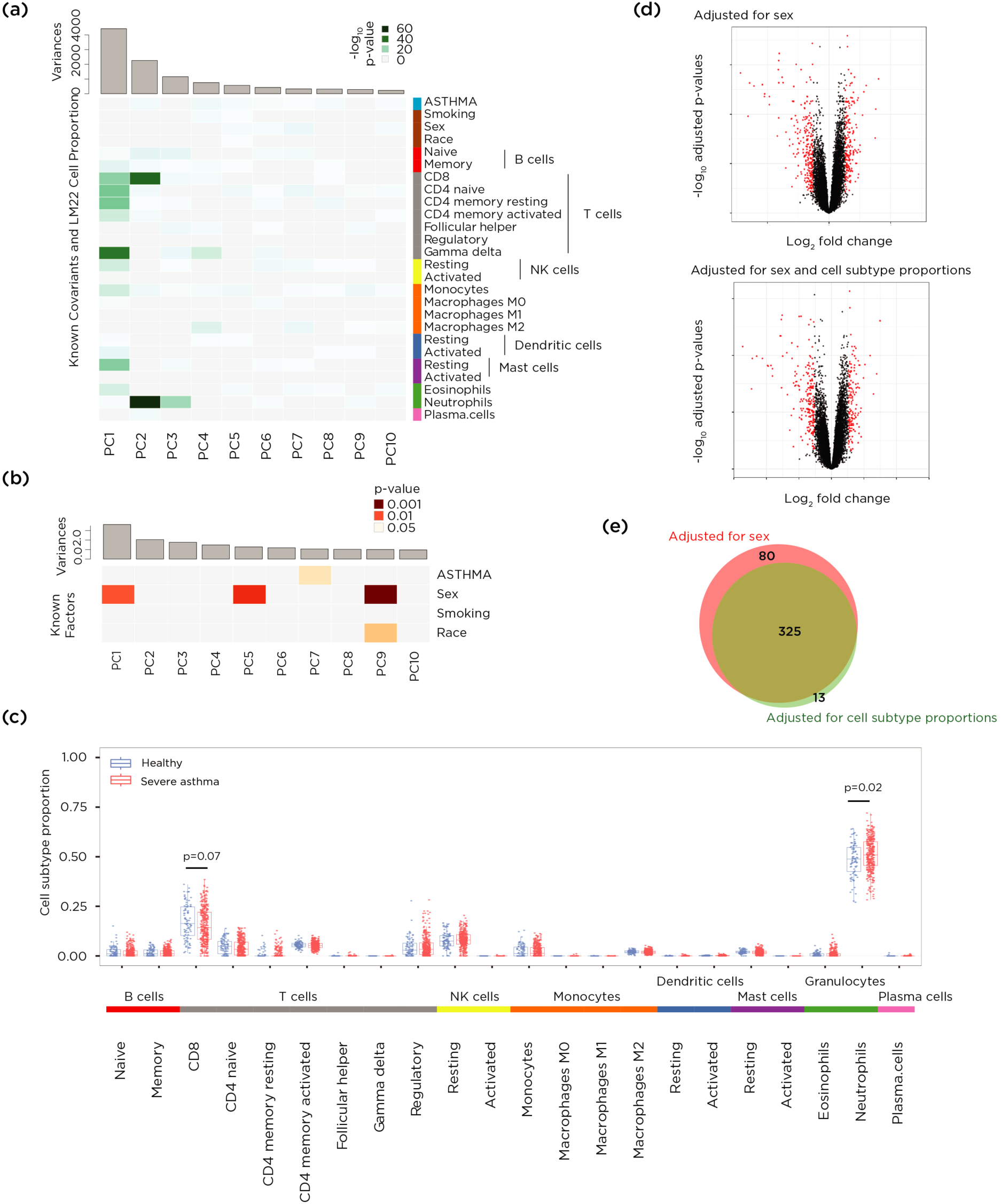
Deconvolution shows a strong effect of cell subtype proportions on gene expression variation in a study of blood leukocytes from asthmatics. (a) Principal components for gene expression with significance of association of different factors shown as a heat map. The disease status of severe asthma (ASTHMA) was very weakly associated with the variability in gene expression, accounting for only 0.31% of the first PC (which accounts for 20.9% of the variance, p=0.25) and 4.1% of the second PC (10.68% of the variance, p=2.9*10^−5^) of expression variation. (b) The same kind of analysis was performed but this time testing the contributions to the cell subtype proportional variability, with small contributions to principal component 1 (accounting for 16.54% of variance) of disease status (0.023%, p=0.75) and sex (1.56%, p=0.01). (c) Looking into why the cell subtypes were so influential in altering gene expression, we find the proportions of two cell types to be significantly different in patients with severe asthma, an increased proportion of neutrophils (p= 0.021, Mann-Whitney test) and a decreased proportion of CD8+ T cells (p=0.072, Mann-Whitney test). In (d) we show two volcano plots, the upper showing 405 differentially-expressed genes (DEGs, FDR-adjusted p value <0.05, >1.2 fold change in expression) in red, representing those identified without cell subtype proportion adjustment. The lower volcano plot shows the 166 DEGs following adjustment for cell subtype proportions using the PCs most strongly reflecting the cell subtype effects on expression variation. (e) The Venn diagram shows the overlap between the genes identified as differentially-expressed when adjusted for sex (red) and after additional adjustment for cell subtype proportional variation (green).

### Cell subtype proportion changes in asthma

We then performed a further PCA to identify the factors in the phenotypic data and metadata that were most associated with the cell subtype proportions observed. The input in this case was the matrix of estimated cell subtype proportions across samples, and the objective was to find the biggest contributors to the variance in cell subtype proportions between samples. We tested these correlations and the significance of the contribution of each phenotypic variable to each PC of the expression profiles. We found no significant correlation with disease status (R^2^=0.00023, p=0.75) and a small but significant contribution of sex (R^2^=0.01557, p=0.011) to the first principal component of the estimated cell subtype proportions (16.54% of variance) (Figure 1b, **Supplementary Table 2**).We observed that the proportion of neutrophils was significantly increased in severe asthma patients compared to healthy controls (p= 0.02, Mann-Whitney test), with a decrease in the proportion of CD8+ T cells (p=0.07, Mann-Whitney test) (Figure 1c). These results are consistent with several prior studies that have reported associations between neutrophils and asthma severity^22–26^. Less is known about the role of CD8+ cells in asthma. While some studies have reported fewer CD8+ T cells in allergic asthma^27^, others have found higher numbers of CD8+ cells in asthmatics that correlated with asthma severity but not with atopy.Since atopy is associated with fewer CD8+ T cells^28^, and most participants in this study were atopic (of the 334 severe asthmatics, atopy information was available for 308, of whom 234 were positive (76.0%); of the 87 controls, atopy information was available for 76, of whom 32 were positive (42.1%)), their atopic status rather than their asthma may explain the reduced CD8+ proportions.Our re-analysis of functional genomics data associates severe asthma in atopic individuals with higher neutrophil and lower CD8+ T cell proportions, which offers potential insights into the cellular events occurring in these individuals. These results suggest that the variation of cell subtype proportions may not be merely a confounding variable in gene expression studies, but can potentially contribute useful insights into the biological processes occurring in a disease.

### Cell-intrinsic gene expression changes in asthma

When the effects of cell subtype proportional changes are eliminated, the changes in gene expression that remain are more likely to represent altered levels of gene transcription within the cells tested. We borrow a term used in the study of DNA methylation to refer to these as cell-intrinsic gene expression changes^5^, reflecting what we have described as cellular reprogramming^13^. Including and adjusting for sex as a covariate in our analyses, we identified 405 differentially-expressed genes (DEGs) without adjusting for cell subtype proportions (FDR-adjusted p-value<0.05, >1.2-fold change in expression, Figure 1d, genes listed in **Supplementary Data 1**).When we performed an adjustment for cell subtype proportions including each of the individual cell proportion values in our linear model, as is typically performed in studies of DNA methylation^17,29,30^, only 142 genes remained categorized as DEGs (listed in **Supplementary Data 2**). However, this approach is not ideal, as it introduces a large number of covariates into a multi-variable linear regression model, and these covariates are collinear with each other (as one cell subtype proportion goes up, other proportions have to go down). We therefore used the alternative approach of regression on PCs of cell subtype proportions^31,32^ using the PCs that most strongly reflected the cell subtype proportion effects on expression variation (those with a p <0.01 and explaining >1% of variation of cell subtype proportions: PCs 1-5 and 9 in **Supplementary Fig. 1**). We thus reduced the dimensions of the covariates and eliminated their collinearity. This PC-based approach now defined 338 genes to be differentially-expressed (Figure 1d, genes listed in **Supplementary Data 3**), eliminating 80 of the 405 DEGs originally identified prior to cell subtype adjustment, and adding a further 13 genes not previously recognized as differentially expressed (Figure 1e).

### Intra-individual cell subtype changes over time

The above example shows that gene expression differences between individuals studied using a cross-sectional design can be influenced by cell subtype proportions. A question that arises is whether these cell subtype proportions are stable within an individual, and the contribution of intra-individual fluctuation of cell subtype proportions to gene expression variation is minimal, or whether these proportions vary between sampling times in the same individual, with substantial effects on gene expression variation. We took advantage of an unusual study that tested peripheral blood mononuclear cells (PBMCs) sampled from the same individual over an 18 month period^20^. We again applied *CIBERSORT* deconvolution and observed substantial variation in cell subtype proportions within this individual over time (**Supplementary Fig. 2**). Once again (after adjustment for RNA integrity number (RIN) variation) we observed a strong cell subtype effect on expression variation, observed in principal components 1-3 and 5 (Figure 2). Based on a multiple linear regression model, the estimated contribution of cell subtype proportions to the principal components of gene expression (PC1 to PC5,>5% proportion variance) was 22.28% (**Supplementary Note 1, section 2-4**). We therefore estimate that intra-individual cell subtype proportion variation may account for more than 20% of the gene expression variation in blood.

**Figure 2.**
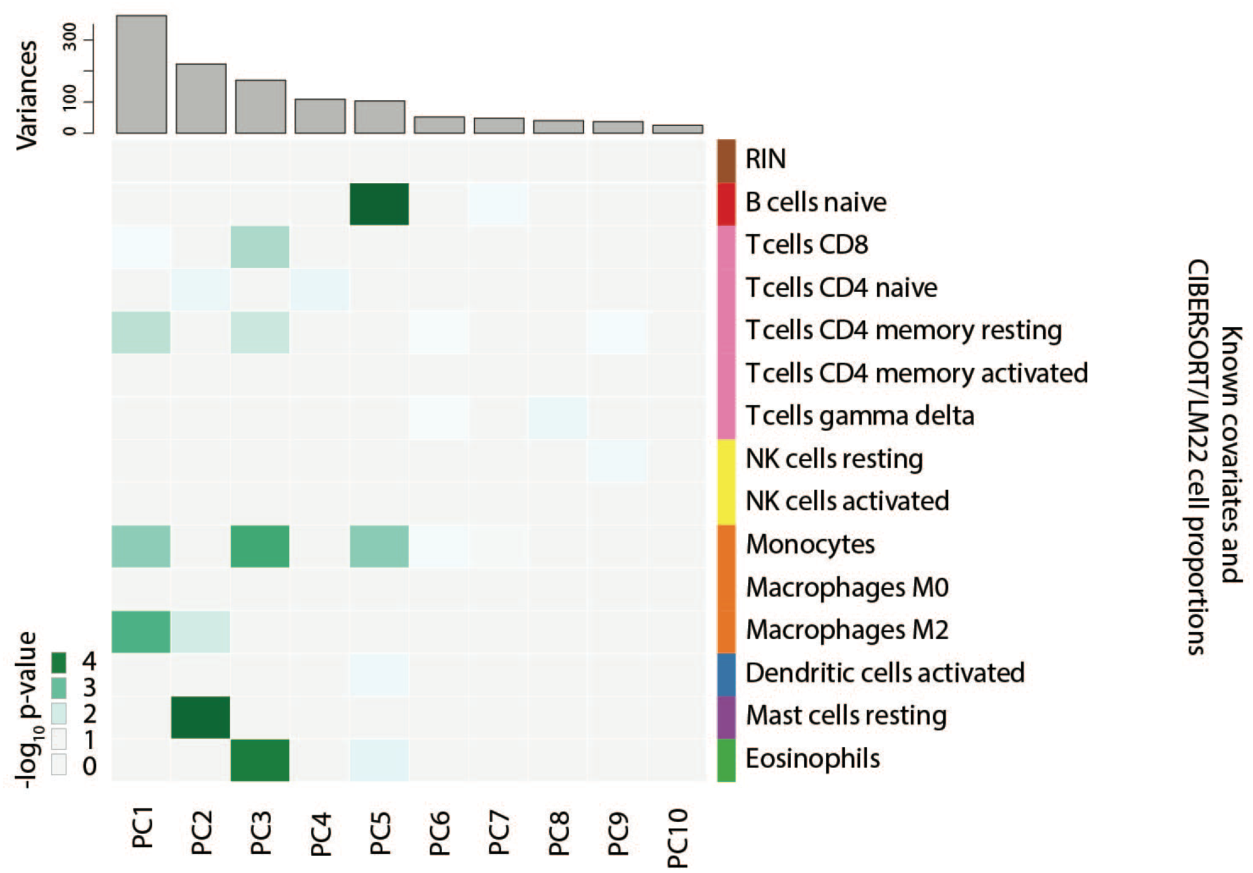
Variable cell subtype proportion profiles were observed in peripheral blood samples collected over 18 months from the same individual. We show a heat map depicting the influences on the variability of gene expression in each principal component. Cell subtype effects are observed in principal components 1-4 and 6 in particular. A multiple linear regression model allows an estimate of the contribution of cell subtype proportion variation to the first principal component (PC) of gene expression which accounts for 24.5% of the variance, finding 53.87% of this PC to be attributable to cell subtypes (model p-value = 0.0083, multiple R^2^=0.5387, adjusted R^2^=0.343).

### Validation of reference-based deconvolution methods

Our next focus was on a study of aging that measured both gene expression and DNA methylation in the same PBMC samples, comparing 146 nonagenarians with 30 young controls^19^. These authors performed flow cytometry to measure the CD4/CD8 ratio of T cell subtypes in these samples, providing a reference against which we could compare our *in silico* predictions of cell subtype proportions. To quantify cell subtypes using DNA methylation data, we used the reference-based approach developed by Houseman and colleagues^1^ that uses reference DNA methylation profiles from 6 blood cell subtypes, including CD4 and CD8 T cells. To generate a CD4+ T cell quantification from the LM22 reference gene expression data^21^, we summed the values for naïve, resting memory and activated CD4+ T cells. We show these results in **Supplementary Fig. 3**.Cell subtype quantification based on DNA methylation estimates CD4/CD8 ratios exceptionally well (R^2^=0.66, root-mean-square error (RMSE)=1.42) while the estimates based on gene expression are also well correlated with the flow cytometry results (R^2^=0.36, RMSE=2.14).

### Concordance of cell subtype predictions by orthogonal assays

We tested the performance of the cell subtype deconvolution approaches a second way, by comparing the concordance of predictions of the proportions of cell subtypes by the orthogonal gene expression and DNA methylation assays performed on the same samples. As the DNA methylation approach tests only 6 cell types^1^, fewer than the 22 tested by gene expression studies^12^, we restricted our comparisons to the 6 cell types in common (**Supplementary Note 2**).We observed a general strong correlation between the estimates of cell subtype proportions based on deconvolution of DNA methylation and gene expression data (r=0.48∼0.77, Figure 3a). We replicated this analysis using data from an independent study that compared 97 atopic asthmatic and 97 non-atopic/non-asthmatic children^17^.Again, we observed similar linear correlations between the estimates based on DNA methylation and gene expression deconvolution (r=0.38∼0.75) (**Supplementary Fig. 4**).

**Figure 3.**
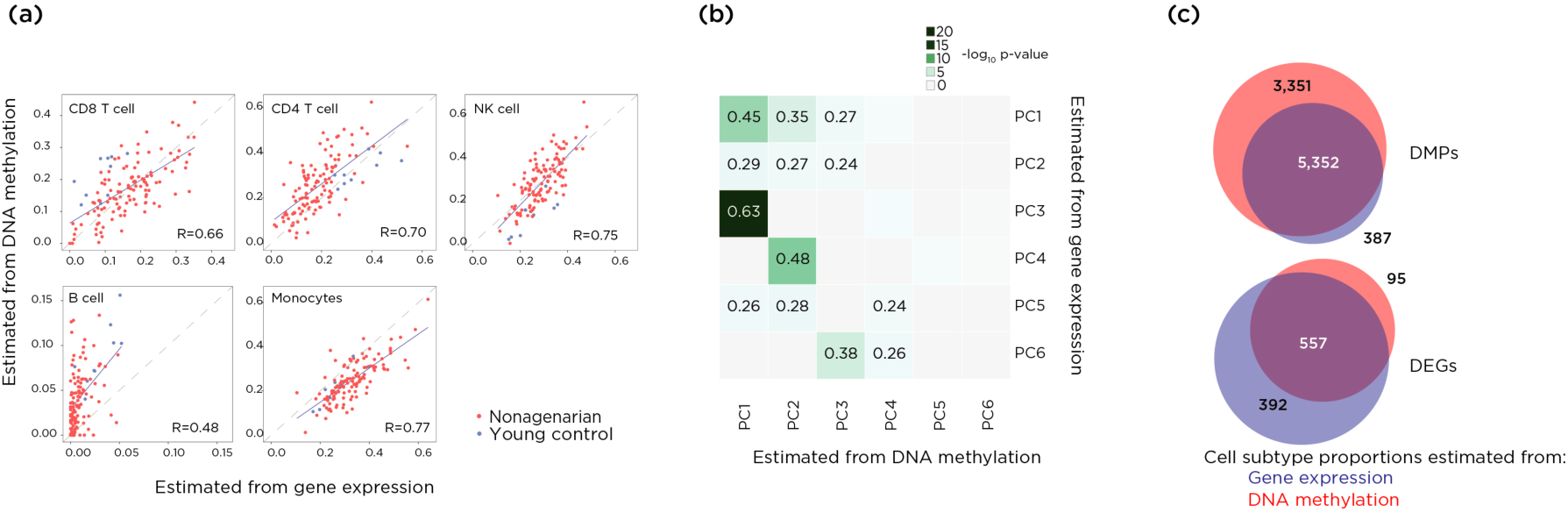
The estimated cell subtype proportions from different genome-wide assays on same samples showed good correlations. (a) The cell subtype proportions estimated by deconvolution of gene expression and DNA methylation data from the same samples are strikingly concordant for most cell types (r=0.48∼0.77). (b) A linear regression modelling approach measured the correlations between the principal components testing cell subtype proportion effects on gene expression and DNA methylation, with strong correlations (r=0.24∼0.63) between the first four PCs.(c) We tested whether adjustment for cell subtype proportions using gene expression and DNA methylation-based deconvolution yielded comparable results. There were 5,739 differentially methylated probes (DMPs) (FDR-adjusted p-values <0.05, beta value differences between groups >10%) after adjusting using gene expression data, and 8,703 after adjusting using DNA methylation data, with the majority (5,352) in common between the two groups. There were 949 DEGs (FDR adjusted p-value<0.05 and >1.5-fold difference between groups) after adjustment for sex and cell subtypes based on gene expression data, and 652 after adjustment based on DNA methylation data, with the large majority (557) of the cell subtype-adjusted DEGs in common between the two groups.

We then explored the correlations between principal components of cell subtype proportions estimated from DNA methylation profiles and principal components of cell composition estimated from gene expression profiles using a linear regression modeling approach. As expected, we observed positive correlations (r=0.24∼0.63) between these principal components (Figure 3b).Next, we tested whether adjusting for cell subtype proportions based on the deconvolution results from gene expression or from DNA methylation assays influenced the same DEGs or differentially-methylated probes (DMPs).We studied 126 individuals on whom both DNA methylation and expression profiles had both been generated as part of the study of aging^19^. Our analysis, without adjusting for cell subtypes, identified 12,309 DMPs (FDR-adjusted pvalues <0.05, beta value differences between groups>10%).After adjusting for cell subtype proportions estimated from DNA methylation data, we identified 8,703 DMPs, and identified 5,739 DMPs after adjusting for cell subtype proportions using gene expression data. Of the 5,739 DMPs retained after the gene expression-based adjustment, 5,352 (93.3%) were concordantly predicted in the group of 8,703 identified after deconvolution using the DNA methylation data (Jaccard similarity index=0.59) (Figure 3c). We performed the same kind of analysis for DEGs, identifying 949 genes as differentially-expressed following deconvolution based on gene expression data (FDR adjusted p-value <0.05 and >1.5-fold difference between groups, Figure 3c), and 652 after adjustment for cell subtypes based on DNA methylation data, with 557 (85.4%) of the 652 DEGs in common (Jaccard similarity index=0.53). The strong concordance of these functional genomics results, whether using gene expression or DNA methylation for deconvolution, indicates that the cell subtype proportions estimated by one assay can be used to test the influence of cell subtype proportional composition in results from a different functional genomic assay performed on the same samples.

### Cell subtype effects are dominant in SLE

Systemic lupus erythematosus (SLE) is an autoimmune disease that involves numerous cell types of the immune system^33^.Our cell subtype deconvolution revealed that the proportion of monocytes was significantly increased and the proportion of resting natural killer (NK) cells was significantly decreased in SLE, getting the same results using either DNA methylation and gene expression cell subtype deconvolution (Figure 4a, **Supplementary Fig. 5a**). These cell subtype proportion changes revealed from functional genomic data are consistent with prior literature describing a lower proportion of NK cells and a higher proportion of monocytes in patients with SLE^34^. In addition, 53.8% of the first principal component of DNA methylation variation (which accounted for 16.3% of variance, p=9.1e^-14^) and 94.9% of the second principal component (10.6% of variance, p=0.084) were attributable to cell subtype variation (Figure 4b). Similar results were obtained from gene expression estimates of cell subtype proportions, with 78.9% of the first (accounting for 12.5% of variance, p=0.012) and 81.8% of the second principal component (9.1% of variance, p=0.005) attributable to cell subtype variation (**Supplementary Fig. 5b)**. When we re-analyzed gene expression differences between SLE and control subjects accounting for cell subtype variability (using gene expression information) we found that only 4 genes of the 485 DEGs (false discovery rate (FDR)<0.05 and log_2_ fold-change (FC)>1.2, **Supplementary Data 4**) identified without adjusting for cell subtype proportions remained significant. In the DNA methylation analysis, we identified 2,154 differentially methylated CGs (FDR <0.05, Δbeta(case-control) ≥10%, at 1,366 genes) without adjusting for cell subtype, but just 40 CGs (at 27 genes) after adjusting for cell subtype proportions (Figure 4c-d, **Supplementary Data 5)**. This suggests that almost all significant differences can be attributed to systematic cell subtype proportion variation between individuals with SLE and healthy controls. When we linked the 40 CGs with the 27 nearby genes (using the Illumina microarray design annotation) and tested gene ontology (GO) terms for enrichment, the list of most significantly enriched GO terms changed compared with the unadjusted data (Figure 4e), revealing new terms including a strong enrichment for type I interferon signaling pathways (Figure 4f). These results clearly indicate that almost all alterations observed between SLE cases and controls samples can be attributed to changes of cell subtype proportions.

**Figure 4.**
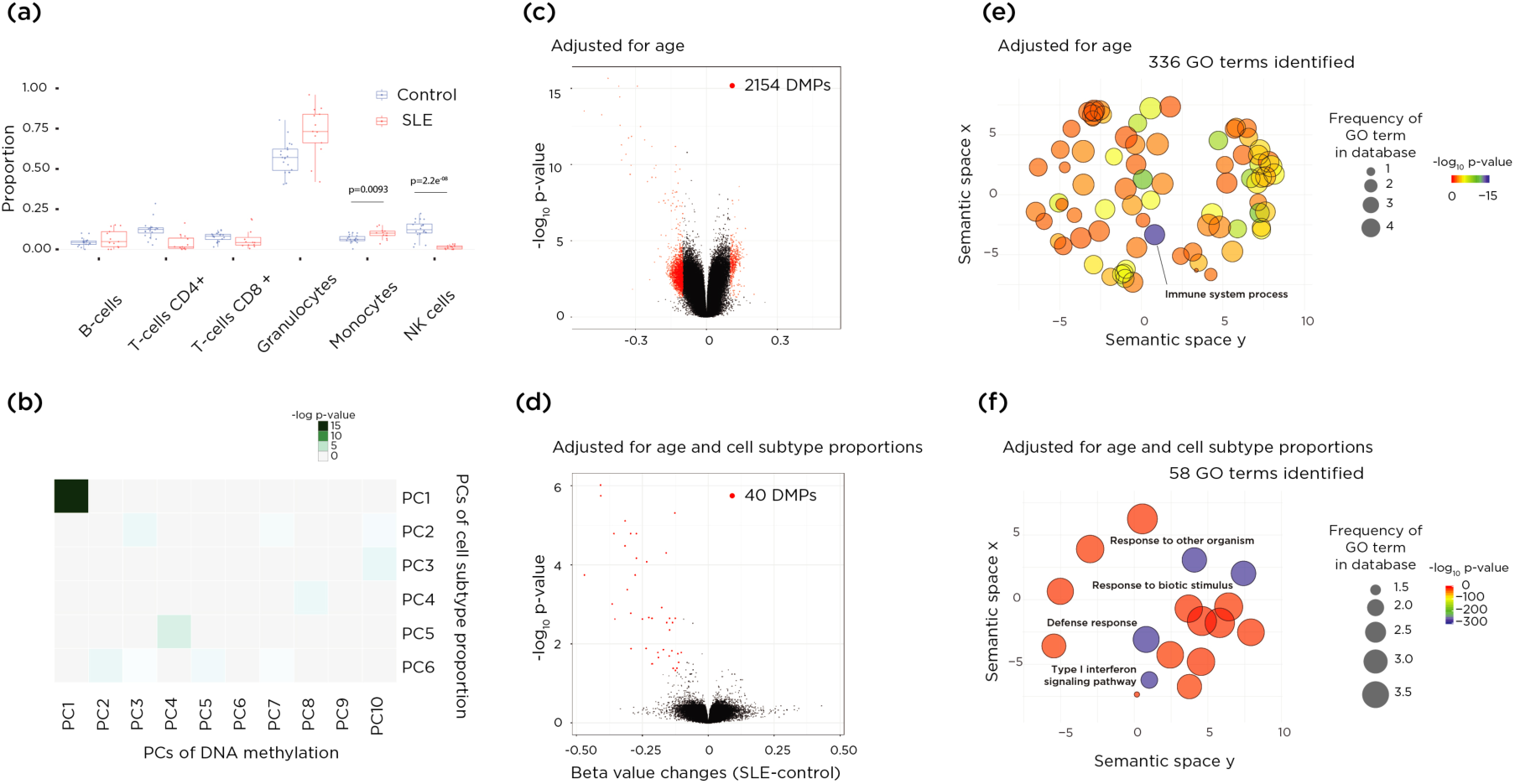
Disease status associated cell subtype proportion changes reflect the DNA methylation changes between SLE and control samples and the disease-related pathways were highlighted after the cell subtype proportion adjustment. (a) The cell subtype proportion changes in SLE were driven predominantly by two cell types. The proportion of monocytes in increased in SLE (p=0.0093, t-test). On the other hand, the proportion of natural killer (NK) cells is higher in patients with SLE (p=2.2e^-08^, t-test). (b) Principal components for DNA methylation with significance of association of PCs for the estimated cell subtype proportions are shown as a heat map. The first principal components were significantly associated each other. (c-d) Two volcano plots show differentially-methylated probes (DMPs). (c) shows the results of DMPs adjusted for age alone and (d) shows results also adjusted for cell subtype proportions. Almost all DMPs were eliminated after the adjustment for cell subtype proportions. (e-f) GO analysis results are summarized as REVIGO scatterplots. (e) shows the results of age-adjusted and (f) the results for age and cell subtype proportion adjusted terms. The x and y axes indicate the semantic similarity of each GO term. The bubble color indicates log_10_ p-values, and size represents the percentage of genes annotated with the GO term in the human database.

### Single-cell RNA-seq data for cell subtype deconvolution

The approaches described above require reference information about gene expression or DNA methylation profiles in the cell subtypes composing the tissue being studied.Such reference profiles are well described for blood, but for most other tissues in the body such reference information will be unavailable. We have proposed that single cell transcription analysis (scRNA-seq) could be used in these situations^13^. The scRNA-seq technique does not require *a priori* insights into cell subtypes present in the tissue studied, and it can be performed on relatively small numbers of cells. To test the potential value of scRNA-seq as a way of generating reference expression data, we downloaded a publically available scRNA-seq data set of ∼68,000 PBMCs (see **Methods**, details of the preprocessing procedures used provided in **Supplementary Note 3**), analyzing these data using *Seurat*^35^. We identified 21 clusters, indicating subtypes of PBMCs, based on expression patterns within each cell (**Supplementary Fig. 6a**).By searching for expression of genes encoding canonical cell subtype markers, we matched the clusters to known PBMC subtypes (**Supplementary Note 3**). We then identified 828 genes, each of which had an expression pattern that was highly cluster-specific and was expressed by at least 50% of cells in the cluster (**Supplementary Data 6**). This allowed us to calculate a median expression value for these cluster-specific signature genes as the basis for a cell subtype signature profile for *CIBERSORT* analysis. We tested the performance of this scRNA-seq reference data set using the CD4:CD8 ratios from the aging study. Using known cell subtype-specific genes, we were able to test for their expression in each cluster (we list these genes in **Supplementary Data 7**). Based on the expression status of those marker genes, we assigned clusters 0, 1, 2, 12 and 13 as CD4+ T cells and clusters 3, 4 and 6 as CD8+ T cells (**Supplementary Data 7**). The result of CD4:CD8 ratio analysis is shown in **Supplementary Fig. 6b.** While the values of R^2^=0.26 and RMSE=2.64 indicate less accurate performance compared with the reference-based approaches (Figure 3a), the scRNA-seq data are relatively predictive of these cell subtype proportions. We also compared how the scRNA-seq approach performed in quantifying proportions of cell subtypes relative to reference-based approaches based on DNA methylation data, finding positive linear correlations (R=0.32 to 0.61) for all cell types except B lymphocytes (**Supplementary Fig. 6c**). We tested the association of PCs of estimated cell subtype proportions using scRNA-seq results with the PCs derived from DNA methylation (Figure 5a) and gene expression (Figure 5b) reference data. We observed comparable correlation to PCs of cell subtype proportion from DNA methylation and LM22. We also tested whether adjusting DEGs and DMPs based on cell subtype proportions inferred using the scRNA-seq signatures gave similar results to the other approaches, and found concordance of the genes and loci identified (Figure 5c).

**Figure 5.**
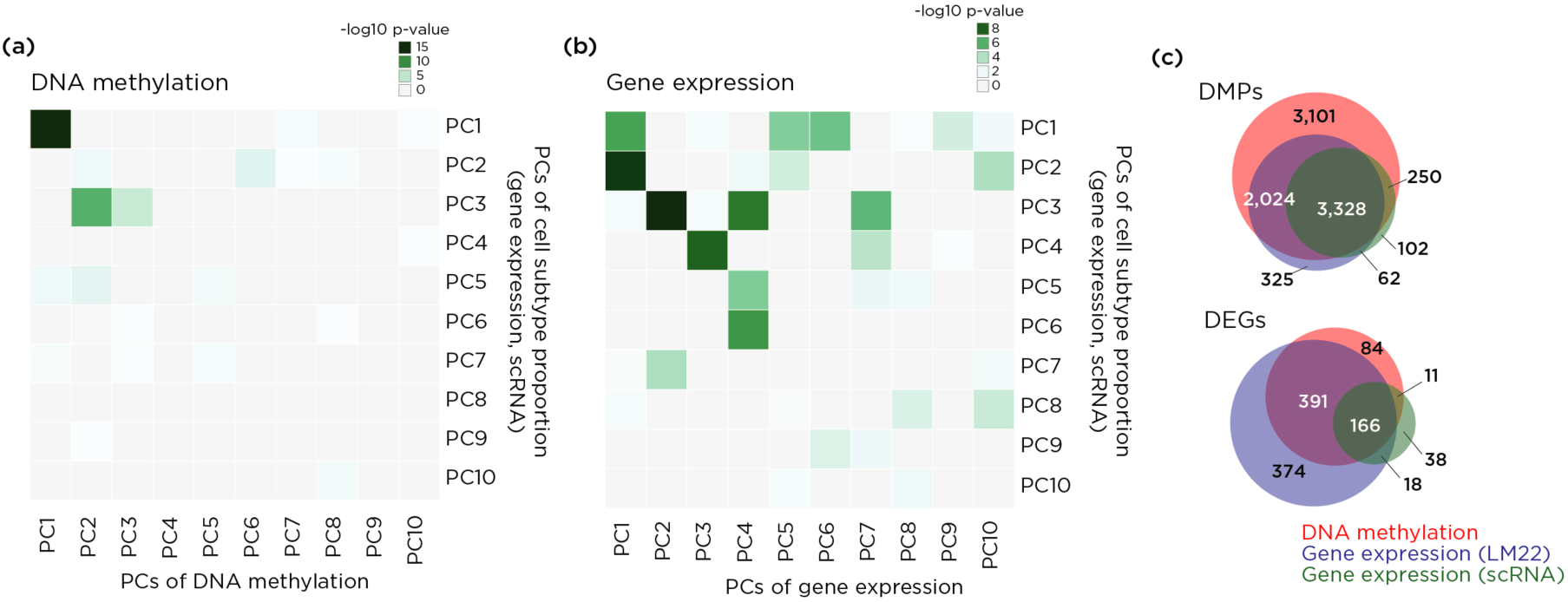
Deconvolution of estimated cell subtype proportions using the signature profile from single-cell RNA-seq (scRNA-seq) results shows comparable results to the estimations using reference profiles from purified cells. A heatmap shows principal components for DNA methylation (a) or gene expression (b) with significance of association of PCs for the estimated cell subtype proportions using scRNA-seq gene signature profile. Panel (c) shows how adjustment of DEGs and DMPs from the aging study based on cell subtype deconvolution using scRNA-seq/*CIBERSORT* generates similar results to the other approaches, showing a strong concordance.

Finally, we tested the stability of the prediction of differentially-methylated positions (DMPs) following the different types of adjustments. These cell subtype proportion adjustments were based on gene expression, DNA methylation or scRNA-seq reference data, either using the cell subtype proportions themselves or the PCs capturing this information, as described earlier. To measure stability, we asked how many DMPs were retained in a list of the same number ranked either by significance (Figure 6a) or by the magnitude in change of beta value (Figure 6b). We find that the 22 cell type gene expression and 6 cell type DNA methylation reference panels work better than the scRNA-seq gene expression reference panel, but that within a reference panel approach the use of PCs is associated with more stable prediction of DMPs than when using unprocessed cell subtype proportion values. In fact, there is a dramatic effect on stability of DMPs ranked by the magnitude of beta-value differences when using the large, collinear models based on cell subtype proportions, as these models grossly alter the beta values, an effect that no longer occurs when using the PC surrogates approach instead (Figure 6b).

**Figure 6.**
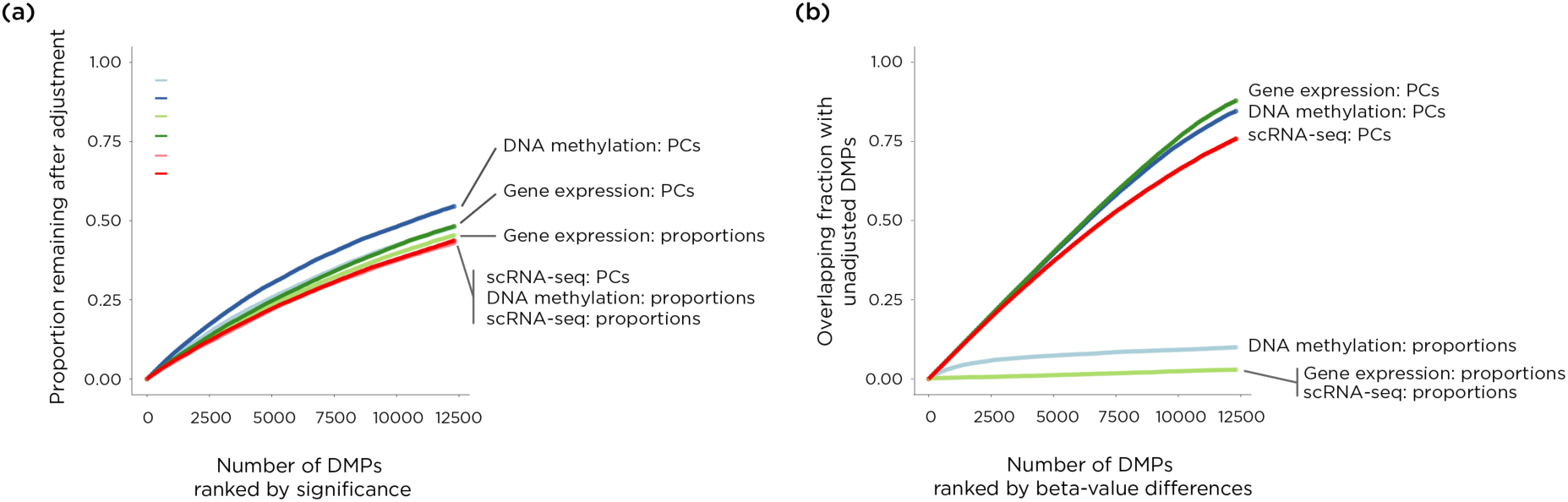
Testing stability of calling differentially methylated probes using different cell subtype proportion adjustment methods shows that PC-based methods improve stability for DMPs ranked either by significance or effect size (adjusted beta-value differences). The plot shows the concordance of the proportion of the top DMPs using the different adjustment methods. We ranked the DMPs either by significance or by degree of difference of beta values, and calculated the overlapping fraction of DMPs prior to adjustment. We show results of stability of DMPs ranked by significance levels (a) and by effect sizes (b).

### Evaluation of a reference-free deconvolution approach

The alternative to reference-based deconvolution is to use a reference-free approach, of which surrogate variable analysis (SVA) is probably the most commonly used in DNA methylation studies^36^. SVA is an attractive approach, as it should allow not only the effects of cell subtype proportion heterogeneity but also influences like batch effects to be eliminated as confounding when looking for cell-intrinsic changes in functional genomics properties. We explored how SVA performs when insights are available from deconvolution into cell subtype composition. We studied the DNA methylation data from the aging study, testing how each surrogate variable was influenced by each of the metadata variables. In Figure 7a we show that SVA does indeed predict cell subtype proportions as surrogate variables, as well as picking up a strong influence of experimental batch. However, despite defining age as the phenotype of interest, we find that the SVA is also recognizing this as a source of variability, which would result in an unrecognized loss of the signal sought in these studies. To simulate a situation in which cell subtype proportion and batch effects are not major confounding influences in an experiment, we re-processed the aging data to remove batch effects on their own or in combination with cell subtype proportion effects.In those situations, the effect of SVA to influence the phenotype of interest gets progressively stronger (Figure 7b-c), disproportionately penalizing what would be better-executed studies.

**Figure 7.**
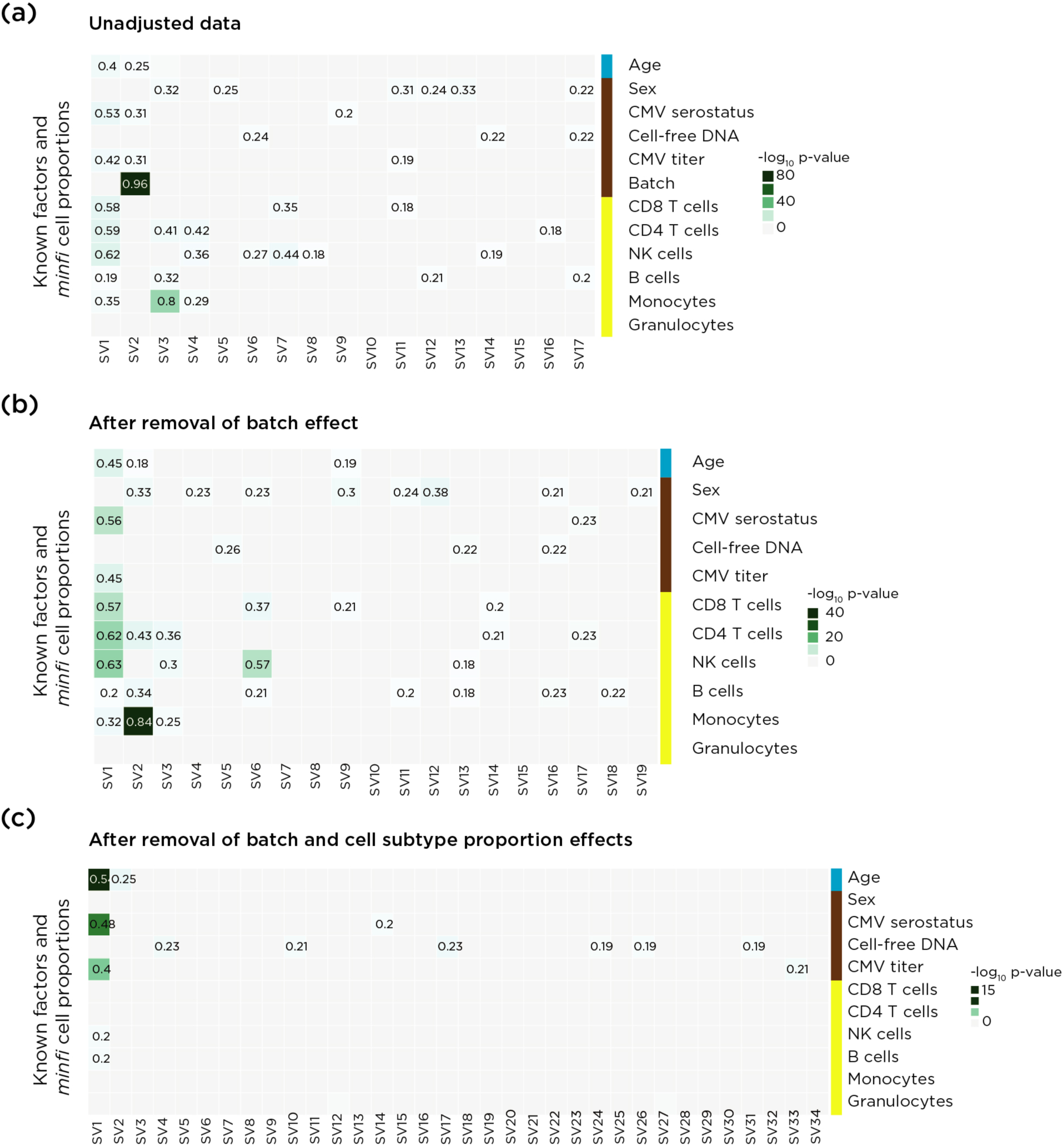
The SVA approach may lead to elimination of true positive results.Heatmaps show the influences on each surrogate variable (SV) of known covariates and estimated cell subtype proportions. We performed SVA on the unadjusted data (a), and on data after removal of batch effect (b) and after further adjustment for cell subtype proportion variability (c). We performed multiple linear regression models to estimate the contribution of covariates to surrogate variables. The step-wise analysis revealed that the SVA approach has effects on the phenotype of interest, especially as we eliminate some of the sources of experimental artefact.

## DISCUSSION

By using assays that test expression of genes or microRNAs, methylation of DNA, chromatin states or other indicators of genomic function, we are generally trying to understand the innate characteristics of the cells tested.Such cell-intrinsic changes can reflect responses to environmental perturbations or genetic mutations, and can be used as clues to the pathogenesis of an associated phenotype. We have referred to this as cellular reprogramming^13^, the alteration of the molecular characteristics of a canonical cell type.The possibility that cell subtype proportional heterogeneity could be contributing to the variability in the results of the functional genomics assay is not always considered, but when addressed is generally treated as a confounding variable with the focus on cell-intrinsic changes of functional genomic properties.

We have pointed out that the systematic alteration of cell fate decisions and the repertoire of cell subtypes in a tissue is a potential outcome of transcriptional regulatory perturbations, potentially contributing to the development of specific phenotypes, an alternative cellular epigenetic model that we have called polycreodism^13^. In the current study, we sought to understand the relative contribution of each cellular epigenetic model, reprogramming and polycreodism, in different types of phenotypes, from physiological studies of aging and serial sampling from a single individual, through the disease phenotypes of asthma and systemic lupus erythematosus. Our focus was on studies of peripheral blood leukocytes, not only because of the technical advantages they offered for our cell subtype deconvolution approaches, but also because many transcriptomic studies and most published large-scale epigenome-wide association studies (EWAS) testing DNA methylation have been performed on blood cells. By gaining insights into the relative contributions of cellular reprogramming and polycreodism in blood cells, we could provide insights into the interpretability of these kinds of prior studies that were focused on testing the cellular reprogramming model alone.

Our studies were based on the ability to estimate cell subtype proportions from gene expression or from DNA methylation data. It was helpful to have an orthogonal quantification of the CD4/CD8 T lymphocyte ratio for comparison in the aging study^19^, allowing us to show that both the gene expression and DNA methylation-based deconvolution approaches were reasonably concordant with these flow cytometry-based results. The concordance of cell subtype proportions in the same samples predicted by the separate gene expression and DNA methylation assays also provided reassurance that these proportions are likely to be reasonably accurate. We note that in tissues other than blood such reference gene expression and DNA methylation data are unlikely to be readily available, prompting us to explore whether scRNA-seq data could be used in these deconvolution studies. While the results were not as accurate as the use of reference data from purified cells, they were positively correlated with the CD4/CD8 ratios and with several of the cell subtype proportions predicted from reference-based deconvolution using DNA methylation data.This strongly suggests that generating and using a scRNA-seq reference for a less well-characterized tissue will be overall a helpful way of understanding the cell subtype proportion source of variation in functional genomic assays of that tissue.

There were other observations made that are of technical importance when performing functional genomics studies. We are concerned that using the SVA approach has the potential to mask some of the genuinely phenotype-associated effects, especially in better-executed studies. We note the strong concordance of results when adjusting for cell subtype proportions using gene expression and DNA methylation data, indicating that deconvolution using results of one functional genomics assay can be used to adjust for cell subtype proportions when analyzing a completely different kind of assay of the same samples. We were also careful to avoid using the individual cell subtype proportions in the multi-variable linear regression model, as they can be numerous and are inherently collinear, instead using regression on PCs^31,32^, choosing the PCs capturing most of the effects of cell subtype variability. These insights should be generally useful when performing and analyzing epigenetic association studies in particular and functional genomic assays in general.

Instead of focusing on the cellular reprogramming model, we generated two outputs from the functional genomics studies. The first was a high-confidence set of genes or loci undergoing alterations in gene expression or DNA methylation, manifesting changes that could not be attributed to cell subtype proportional variability, indicating cellular reprogramming effects. The second was the difference in cell subtype proportions between the comparison groups. This is not typically an output of analytical approaches used for gene expression or DNA methylation studies, but was an explicit output of our analytical approach, and revealed systematic changes.It should be relatively straightforward to modify excellent software packages such as *minfi*^37^ to allow this additional output to be generated routinely. In particular, the study of SLE was striking for having an overwhelming effect of cell subtypes on gene expression and DNA methylation variation. While this might currently be considered a negative result, treating cell subtype effects purely as a confounding variable, we note that the SLE patients had distinctive NK cell and monocyte proportions, which represents the use of functional genomic data to gain an insight into cellular events contributing to the disease process. These cellular changes had already been recognized independently in SLE, with decreased NK cell activity correlating with active disease and observed to a greater extent among those with renal involvement^38–41^. Conversely, more activated monocytes have been found in individuals with SLE^42,43^, which are associated with disease complications such as atherosclerosis in these patients^44^. This SLE example represents the value of looking simultaneously for cellular reprogramming and cell repertoire changes in functional genomics studies, as each can be harvested from the functional genomics data generated and can be valuable in providing insights into the condition being studied. Reference-free approaches, on the other hand, will eliminate this useful information, another reason for caution in choosing approaches such as SVA.

We conclude that, while it should not be surprising that cell subtype effects need to be taken into account in the interpretation of functional genomics studies, it is probably worth paying more attention to the characterization of the variability of cell subtypes as an insight into the phenotype being tested, rather than discarding the information as merely confounding^1,45–47^. Phenotypes may indeed result from cellular reprogramming, but it is highly plausible that the polycreodism model of altered cell repertoires in a tissue is another potentially very powerful mechanism for mediation of phenotypic changes.By testing simultaneously for the cellular models of reprogramming and polycreodism, we increase our capacity for discovery of new insights into pathogenesis of diseases or the development of other phenotypes.

## METHODS

### Dataset used in this study and preprocessing data

All datasets used in this study are published and publically available through the Gene Expression Omnibus (GEO, https://www.ncbi.nlm.nih.gov/geo), from which we downloaded the datasets.The GEO accession numbers and study designs are described in Table 1. Phenotypic data were extracted from the matrix tables provided by the authors on the GEO website. Before our reanalyses, we tested the quality of data, including batch effects and possible sample swapping, excluding samples when the information about sex provided by the authors was discordant with the data obtained from the sex chromosomes. For the DNA methylation datasets, we first filtered out poor quality samples by testing detection p-value distributions to see background noise level and eliminating samples with high background (average detection p-values>0.01), and by performing PCA, which found a single sample to cluster very distinctly from all of the others, causing us to remove it from further analysis. We then performed quantile normalization using the *preprocessQuantile()* function in the *minfi* R package^37^, and filtering out probes (a) have failed to hybridize (detection p-values >0.01), (b) probes overlapping with and around known SNPs and 1000G SNPs (MAF>0.1), (c) probes that have been shown to be cross-reactive^48^, and (d) probes on sex chromosomes (except study 4, for which we only used female samples). For expression datasets, we aggregated each transcript value by HUGO gene symbol to calculate mean expression values for each gene. The details of the studies and the preprocessing procedures are provided in the **Supplementary Note 1**.

### Reference-based estimation of cell subtype proportions

We estimated cell subtype proportions based on gene expression and DNA methylation data.From the DNA methylation profiles, we estimated the proportions of CD8+ T cells, CD4+ T cells, NK cells, B cells, monocytes and granulocytes using *the estimateCellCounts()* function from the *minfi* R package^37^, which is modified from the original Houseman reference-based approach^1^. From gene expression profiles, we ran *CIBERSORT*^12^ using two different signature gene files, the *CIBERSORT* default file based on expression profiles from 22 leukocyte subtypes (LM22)^21^, and a signature gene profile generated from publicly-available scRNA-seq results from 68,000 PBMCs^49^ using the *Seurat* R package^35^.

### Associations between cell subtype proportions and phenotype

We performed principal component analysis (PCA) on the cell subtype proportion estimates obtained. We tested for possible confounding influences using metadata provided by the study authors as a matrix table, including technical (RIN, sample collection date) and biological (age, sex, phenotype) influences, using a linear modeling approach.We identified significant confounding covariates using ANOVA.

### Contribution of cell subtype proportion to functional genomic data

We performed PCA on gene expression values (aggregated expression values) and DNA methylation values (quantile normalized M values), then we tested the contribution of cell subtype proportions to each principal component (PC) using a linear modeling approach. The degree of contribution to each PC was estimated by the R-squared of the regression model and the significance of each was tested using ANOVA.

### Identifying gene expression and DNA methylation changes

To identify differentially methylated probes (DMPs) and differentially expressed genes (DEGs), M values of DNA methylation data and log-transformed values of expression data were used in the regression models with the *lmFit* function of the *limma* R package^50^.We selected biological covariates provided by the authors to be included into the model based on data from each PCA (the covariates used for each study are described in the **Supplementary Note 1**).We built models with and without cell subtype proportion adjustments to test the effects of variability of cell subtype proportions. To avoid collinearity and high dimensionality of cell subtype estimates^51^, we used a principal component regression instead of a linear regression approach using the actual cell proportions. The PCs we included in the linear model are those with significant associations with DNA methylation or expression variation (p-value <0.01) and which explain >1% of the variation of the cell subtype estimate. To identify significant DMPs, we retained the CpGs with FDR<0.05 and absolute beta value changes >10%. The DEGs were defined as the genes with FDR <0.05 and absolute fold changes of expression for studies 1-4 of >log_2_(1.2) and >log_2_(1.5) for study 5 using the same fold-difference threshold as the original publications. The proportional Venn diagrams were plotted using BioVenn^52^.

### Gene ontology analyses

To identify the enriched gene ontology (GO) terms in the DMPs, we performed GO analysis using the Bioconductor package *GOseq*^1^. We used DMP corresponding gene symbols for searching enriched GO terms in the human hg19 database. We selected the terms which false discovery rate (FDR) adjusted p-values were less than 5% as significant GO terms. We performed the analysis on both with and without adjusting for cell subtype proportions. The significant GO terms were visualized using REVIGO^1^, using the program’s default settings (*Homo sapiens* database).

### Single cell RNA-seq analyses

We downloaded the 68,000 PBMC scRNA-seq data set from the 10X Genomics website^49^ and analyzed the scRNA-seq data using the *Seurat* R package^35^. The data contains 32,738 genes in 68,579 cells, with a median number of detected genes per cell of 525 (range 153 to 2,740). After filtering out the genes with fewer than 3 cells expressing the gene and cells in which fewer than 200 genes were found to be expressed, we were left with 17,787 genes in 68,262 cells. The median percent of mitochondrial DNA gene representation was 0.016.After adjusting for the number of unique identifiers per cell and the percent of mitochondrial genes, we performed PCA for linear dimensional reduction. We identified 21 clusters in total, corresponding to 828 signature genes with distinctive expression status compared to other clusters, with on average at least 2-fold differences between the cluster compared with other clusters, and with at least 50% of the cells in the cluster expressing the gene using the *FindAllMarkers* function of the *Seurat* R package^35^. We calculated the median expression values of the signature genes in each cluster to generate a cell subtype signature profile for *CIBERSORT* analysis.We provide the list of signature genes in **Supplementary Data 6** and candidate cell types of each cluster based on the expression status of known genes in **Supplementary Data 7**.

### Surrogate variable analysis

We performed surrogate variable analysis (SVA) using the R package *sva*^14,15^. We selected the phenotype of interest information (young control or nonagenarians) for the analysis. We obtained 17 surrogate variables (SVs) on raw data, 19 SVs after the batch effect adjustment, and 34 SVs after adjustment for batch and cell subtype proportion effects (**Supplementary Note 4**).We tested the correlations to known and estimated cell subtype proportions using a mixed linear regression analysis.We included the SVs in the linear model to test the effects on DNA methylation status.To identify significant DMPs, we retained the CpGs with FDR <0.05 and absolute beta value changes >10%.

### Data availability

All datasets used on this study were downloaded from the Gene Expression Omnibus (GEO, https://www.ncbi.nlm.nih.gov/geo, Table 1). All the customized code used in this study are publicly available at our GitHub server: https://github.com/GreallyLab/PBMC_Kong_2017

## Acknowledgements

We thank Dr. Fabien Delahaye for providing a list of Illumina 450k array probes overlapping with and around known SNPs and 1000G SNPs (MAF >0.1).

## Author contributions

Wrote manuscript draft (YK, DR, CS, JMG, MS), prepared illustrations (YK, JMG, MS), approved final manuscript (YK, DR, CS, JMG, MS), performed experiments (YK, MS), conceived project (YK, JMG, MS), analyzed data (YK, MS), designed experiments (YK, JMG, MS), formulated research questions (YK, DR, CS, JMG, MS), interpreted results (YK, DR, CS, JMG, MS), led investigation (MS).

## Materials and correspondence

Correspondence and materials requests should be addressed to Masako Suzuki.

## REFERENCES

1. Houseman, E. A., et al. DNA methylation arrays as surrogate measures of cell mixture distribution. BMC Bioinformatics 13, 86 (2012).

2. Houseman, E. A., et al. Reference-free deconvolution of DNA methylation data and mediation by cell composition effects. BMC Bioinformatics 17, 259 (2016).

3. Zou, J., Lippert, C., Heckerman, D., Aryee, M., & Listgarten, J., Epigenome-wide association studies without the need for cell-type composition. Nat Methods 11, 309–311 (311).

4. Rahmani, E., et al. Sparse PCA corrects for cell type heterogeneity in epigenome-wide association studies. Nat Methods 13, 443–445 (445).

5. Horvath, S., et al. An epigenetic clock analysis of race/ethnicity, sex, and coronary heart disease. Genome Biol 17, 171 (2016).

6. Shen-Orr, S. S., et al. Cell type-specific gene expression differences in complex tissues. Nat Methods 7, 287–289 (289).

7. Lu, P., Nakorchevski, y. A., & Marcotte, E. M. Expression deconvolution: a reinterpretation of DNA microarray data reveals dynamic changes in cell populations. Proc Natl Acad Sci U S A 100, 10370–10375 (10375).

8. Wang, M., Master, S. R., & Chodosh, L. A. Computational expression deconvolution in a complex mammalian organ. BMC Bioinformatics 7, 328 (2006).

9. Chikina, M., Zaslavsk, y. E., & Sealfon, S. C. CellCODE: a robust latent variable approach to differential expression analysis for heterogeneous cell populations. Bioinformatics 31, 1584–1591 (1591).

10. Gaujoux, R. & Seoighe, C. CellMix: a comprehensive toolbox for gene expression deconvolution. Bioinformatics 29, 2211–2212 (2212).

11. Gon, g. T., & Szustakowski, J. D. DeconRNASeq: a statistical framework for deconvolution of heterogeneous tissue samples based on mRNA-Seq data. Bioinformatics 29, 1083–1085 (1085).

12. Newman, A. M., et al. Robust enumeration of cell subsets from tissue expression profiles. Nat Methods 12, 453–457 (457).

13. Lappalainen, T., & Greally, J. M. Associating cellular epigenetic models with human phenotypes. Nat Rev Genet 18, 441–451 (451).

14. Leek, J. T., & Storey, J. D. Capturing heterogeneity in gene expression studies by surrogate variable analysis. PLoS Genet 3, 1724–1735 (1735).

15. Leek, J. T., Johnson, W. E., Parker, H. S., Jaffe, A. E., & Storey, J. D. The sva package for removing batch effects and other unwanted variation in high-throughput experiments. Bioinformatics 28, 882–883 (883).

16. Bigler, J., et al. A Severe Asthma Disease Signature from Gene Expression Profiling of Peripheral Blood from U-BIOPRED Cohorts. Am J Respir Crit Care Med 195, 1311–1320 (1320).

17. Yang, I. V., et al. DNA methylation and childhood asthma in the inner city. J Allergy Clin Immunol 136, 69–80 (80).

18. Zhu, H., et al. Whole-genome transcription and DNA methylation analysis of peripheral blood mononuclear cells identified aberrant gene regulation pathways in systemic lupus erythematosus. Arthritis Res Ther 18, 162 (2016).

19. Marttila, S., et al. Transcriptional analysis reveals gender-specific changes in the aging of the human immune system. PLoS ONE 8, e66229 (2013).

20. Poldrack, R. A., et al. Long-term neural and physiological phenotyping of a single human. Nat Commun 6, 8885 (2015).

21. Abbas, A. R., Wolslegel, K., Seshasayee, D., Modrusan, Z., & Clark, H. F. Deconvolution of blood microarray data identifies cellular activation patterns in systemic lupus erythematosus. PLoS ONE 4, e6098 (2009).

22. Uddin, M., et al. Prosurvival activity for airway neutrophils in severe asthma. Thorax 65, 684–689 (2010).

23. Moore, W. C., et al. Sputum neutrophil counts are associated with more severe asthma phenotypes using cluster analysis. J Allergy Clin Immunol 133, 1557–63.e5 (2014).

24. Mann, B. S., & Chung, K. F. Blood neutrophil activation markers in severe asthma: lack of inhibition by prednisolone therapy. Respir Res 7, 59 (2006).

25. Kikuchi, S., Nagata, M., Kikuchi, I., Hagiwara, K., & Kanazawa, M. Association between neutrophilic and eosinophilic inflammation in patients with severe persistent asthma. Int Arch Allergy Immunol 137 Suppl 1, 7–11 (11).

26. Cundall, M., et al. Neutrophil-derived matrix metalloproteinase-9 is increased in severe asthma and poorly inhibited by glucocorticoids. J Allergy Clin Immunol 112, 1064–1071 (1071).

27. Tsoumakidou, M., Tzanakis, N., Kyriakou, D., Chrysofaki, s. G., & Siafakas, N. M. Inflammatory cell profiles and T-lymphocyte subsets in chronic obstructive pulmonary disease and severe persistent asthma. Clin Exp Allergy 34, 234–240 (240).

28. Betts, R. J., & Kemeny, D. M. CD8+ T cells in asthma: friend or foe? Pharmacol Ther 121, 123–131 (2009).

29. Steegenga, W. T., et al. Genome-wide age-related changes in DNA methylation and gene expression in human PBMCs. Age (Dordr) 36, 9648 (2014).

30. Pfeiffer, L., et al. DNA methylation of lipid-related genes affects blood lipid levels. Circ Cardiovasc Genet 8, 334–342 (342).

31. Wang, K., & Abbott, D. A principal components regression approach to multilocus genetic association studies. Genet Epidemiol 32, 108–118 (118).

32. Jolliffe, I. T. A note on the use of principal components in regression. Appl Stat 31, 300 (1982).

33. Mok, C. C., & Lau, C. S. Pathogenesis of systemic lupus erythematosus. J Clin Pathol 56, 481–490 (2003).

34. Smith, E., et al. Cross-talk between iNKT cells and monocytes triggers an atheroprotective immune response in SLE patients with asymptomatic plaque. Sci Immunol 1, (2016).

35. Satija, R., Farrell, J. A., Gennert, D., Schier, A. F., & Regev, A. Spatial reconstruction of single-cell gene expression data. Nat Biotechnol 33, 495–502 (502).

36. Teschendorff, A. E., & Relton, C. L. Statistical and integrative system-level analysis of DNA methylation data. Nat Rev Genet (2017). doi:10.1038/nrg.2017.86

37. Aryee, M. J., et al. Minfi: a flexible and comprehensive Bioconductor package for the analysis of Infinium DNA methylation microarrays. Bioinformatics 30, 1363–1369 (1369).

38. Park, Y.-W. et al. Impaired differentiation and cytotoxicity of natural killer cells in systemic lupus erythematosus. Arthritis Rheum 60, 1753–1763 (1763).

39. Green, M. R. J. et al. Natural killer cell activity in families of patients with systemic lupus erythematosus: demonstration of a killing defect in patients. Clin Exp Immunol 141, 165–173 (173).

40. Haga, H. J., et al. Calprotectin in patients with systemic lupus erythematosus: relation to clinical and laboratory parameters of disease activity. Lupus 2, 47–50 (50).

41. Biesen, R., et al. Sialic acid-binding Ig-like lectin 1 expression in inflammatory and resident monocytes is a potential biomarker for monitoring disease activity and success of therapy in systemic lupus erythematosus. Arthritis Rheum 58, 1136–1145 (1145).

42. Henriques, A., et al. Functional characterization of peripheral blood dendritic cells and monocytes in systemic lupus erythematosus. Rheumatol Int 32, 863–869 (869).

43. Byrne, J. C., et al. Genetics of SLE: functional relevance for monocytes/macrophages in disease. Clin Dev Immunol 2012, 582352 (2012).

44. Mikołajczyk, T. P., et al. Heterogeneity of peripheral blood monocytes, endothelial dysfunction and subclinical atherosclerosis in patients with systemic lupus erythematosus. Lupus 25, 18–27 (27).

45. Chen, W., et al. An epigenome-wide association study of total serum IgE in Hispanic children. J Allergy Clin Immunol 140, 571–577 (577).

46. Kinoshita, M., et al. Aberrant DNA methylation of blood in schizophrenia by adjusting for estimated cellular proportions. Neuromolecular Med 16, 697–703 (703).

47. Soriano-Tárraga, C., et al. Epigenome-wide association study identifies TXNIP gene associated with type 2 diabetes mellitus and sustained hyperglycemia. Hum Mol Genet 25, 609–619 (619).

48. Chen, Y., et al. Discovery of cross-reactive probes and polymorphic CpGs in the Illumina Infinium HumanMethylation450 microarray. Epigenetics 8, 203–209 (209).

49. Zheng, G. X. Y. et al. Massively parallel digital transcriptional profiling of single cells. Nat Commun 8, 14049 (2017).

50. Ritchie, M. E., et al. limma powers differential expression analyses for RNA-sequencing and microarray studies. Nucleic Acids Res 43, e47 (2015).

51. Farrar, D. E., & Glauber, R. R. Multicollinearity in Regression Analysis: The Problem Revisited. Rev Econ Stat 49, 92 (1967).

52. Hulsen, T., de Vlie, g. J., & Alkema, W. BioVenn – a web application for the comparison and visualization of biological lists using area-proportional Venn diagrams. BMC Genomics 9, 488 (2008).

